# Geographic Confounding in Genome-Wide Association Studies

**DOI:** 10.1101/2021.03.18.435971

**Authors:** Abdel Abdellaoui, Karin J.H. Verweij, Michel G. Nivard

## Abstract

Gene-environment correlations can bias associations between genetic variants and complex traits in genome-wide association studies (GWASs). Here, we control for geographic sources of gene-environment correlation in GWASs on 56 complex traits (N=69,772–271,457). Controlling for geographic region significantly decreases heritability signals for SES-related traits, most strongly for educational attainment and income, indicating that socio-economic differences between regions induce gene-environment correlations that become part of the polygenic signal. For most other complex traits investigated, genetic correlations with educational attainment and income are significantly reduced, most significantly for traits related to BMI, sedentary behavior, and substance use. Controlling for current address has greater impact on the polygenic signal than birth place, suggesting both active and passive sources of gene-environment correlations. Our results show that societal sources of social stratification that extend beyond families introduce regional-level gene-environment correlations that affect GWAS results.

## Introduction

Genome-wide association studies (GWASs) are an important tool for the investigation of the epidemiology and biology of mental and physical health outcomes. GWASs are viewed as essential herein, because individual differences in almost all outcomes in life are a consequence of the effects of a multitude of genetic and environmental influences.^1^ The aim of a GWAS is to estimate associations between common genetic variants and complex traits.^2^ GWASs set out to estimate the effect of substituting one allele for an alternate allele on the value of a trait or probability of a disease, usually for millions of genome-wide loci. As actual physical substitution of alleles in an experimental setup in humans is ethically undesirable and methodologically challenging, the associations of naturally occurring common genetic variations with naturally occurring trait or disease variations are estimated. One of the underlying assumptions of a GWAS is therefore that estimated relationship between an allele and an outcome is reflective of a causal effect of that allele on the outcome. For the estimated observational effect and the underlying causal effect to be equal, requires the absence of, or adequate statistical control for confounding factors that may introduce allele-trait associations. Confounding is often a result of the presence of unmodeled gene-environment correlation. Three major sources of gene-environment correlation are: 1) population stratification, 2) familial confounding, and 3) confounding due to regional differences in socio-economic factors, which we will refer to as geographic confounding.

Confounding due to population stratification can occur when cultural, behavioral, disease, or other characteristic differences between groups with different ancestries co-occur with allele frequency differences between the ancestries.^3,4^ Allele frequency differences between populations are readily introduced by genetic drift and/or natural selection. Even within reasonably homogenous populations these effects can manifest.^5–7^ These ancestry differences generally show strong correlations with geography,^5–8^ which can result in correlations with environmental influences that show regional differences. To reduce biased genetic estimates in GWASs due to population stratification, it is common practice to run analyses in one of the major ancestral groups (e.g., Europeans, Africans, Asians) and to -within this group-compute principal components (PCs) that reflect the strongest axes of (ancestral) genetic variation and include these as covariates in GWASs.^9^ The efficacy of this method in reducing biases due to population stratification has been evaluated with LD score regression, a method that enables differentiation between confounding (cryptic relatedness and population stratification) from polygenicity in GWASs.^10^ This work revealed that correction for PCs greatly reduced, but did not always fully remove, confounding introduced by ancestral confounding. Recent work has shown that residual traces of ancestral confounding can bias parameter estimates in analyses that use genome-wide summary data to infer the presence or absence of (negative) selection on a trait in recent history.^11,12^

A second process that introduces gene-environment correlations and may bias parameter estimates in GWASs is familial confounding. Here, the parental genotype influences a child’s outcome via the rearing environment (indirect genetic effect), while half of those genotypes are transmitted to the child asserting a direct genetic effect on the outcome. The presence of indirect genetic effects has been demonstrated for genes associated with educational attainment on a variety of outcomes.^13^ When not accounted for, familial confounding will bias genetic effect estimates in GWASs. Moreover, familial confounding is not detectable in LD score regression analysis, as the indirect genetic effect is a true genetic effect, albeit not directly through the effect of the child’s genotype on the trait, but indirectly via its rearing environment. A point worth making is that the precise mechanism underlying the indirect genetic effect remains unclear. Some refer to it as “nature of nurture”^13^, implying indirect effect arises by ways of parenting or nurturing a child. Others call it “dynastic effects”^14^, suggesting that indirect effects can arise, for example, from the succession of (economic) (dis)advantage accumulated by parents to their children, improving their child’s (socio-economic) position at birth. It is important to realize that while most statistical designs rely on family data to infer the presence of an indirect genetic effect, this does not mean that all indirect genetic effects arise specifically due to parental influences. As families are nested in neighborhoods, regions, and other social structures, accounting for family level effects may partly account for effects that take place at various other levels of analysis as well.

A third source of confounding is gene-environment correlations due to regional differences in, often socio-economic, environmental factors, which we, from hereon, will refer to as geographic confounding. Geographic confounding can entail both active and passive processes. Active processes include for example selective migration or brain drain.^15^ Selective migration is a form of active gene-environment correlation, where individuals with favorable genetic predispositions can leverage their skills to improve their environmental circumstances by moving to an economically/socially more favorable region, which in turn improves their outcomes in life. Migration influenced by genotype may improve the rearing environment for offspring as well, and thus socioeconomic confounding through active gene-environment correlation in one generation may induce familial confounding in the next generation. Passive sources of regional gene-environment correlation could be induced by government policies that affect certain socio-economic strata of the population more than others; when the affected groups have different genetic predispositions than non-affected groups, this can introduce a correlation between genotype and (un)favorable environments. For instance, a policy change that makes insulin more expensive and therefore more difficult to purchase for low income groups, will introduce a correlation between alleles related to educational success (which relate to income and SES) and those related to consequences of untreated diabetes. Our analyses are designed to detect the effect of active and passive gene-environment correlation, but for various reasons we outline in the discussion, our results require careful interpretation.

We argue that current GWAS designs generally adequately (but perhaps not fully) deal with population stratification and that designs exist to detect and correct for familial confounding as well.^13,16,17^ However, geographic confounding has been shown to exist,^15^ and current GWAS approaches do not model for it adequately. We further argue that active gene-environment correlation driven by socio-economic processes could have consequences for the way in which genetic associations are interpreted, even when these estimates are obtained from more sophisticated within-family GWAS designs.

In this study, we will investigate the effects of geographic confounding on polygenic signals for a wide range of traits. We will conduct GWASs on 56 complex traits in a dataset of up to 271,457 adult individuals of European descent from Great Britain (UK Biobank)^18^. In these GWASs, we will reduce confounding effects by introducing fixed effects for neighborhoods which are socially and economically more homogenous than a country or study population as a whole. By introducing region fixed effects for relatively homogenous regions into a GWAS, we perform within region GWASs for a wide range of complex traits. We will attempt to distinguish between passive and active processes by basing the regional information on birth place and current address, respectively. We will investigate the impact of passive and active gene-environment correlation on the genome-wide signal of complex traits, as well as on their genetic relationships with socio-economic outcomes (educational attainment and household income).

## Results

### Data and Analysis

We ran linear-mixed model (LMM) GWASs on 1,246,531 single nucleotide polymorphisms (SNPs) for 56 traits with sample sizes ranging from 69,772 to 271,457 UK Biobank participants of European descent. The LMM GWAS controls for cryptic relatedness and population stratification by including a genetic relatedness matrix (GRM) in the model.^19^ As an additional control for population stratification, we included the first 100 PCs derived from the GRM. We also control for sex and age. The GWASs were run on 56 complex traits related to physical and mental health, body composition, and emotional, cognitive, behavioral, and socio-economic outcomes (see Supplementary Table 1 for full list of traits and sample sizes). In order to control for geographic confounding, we included the region of birth and/or current residence of the participants as fixed effects dummy variables. The regions were obtained by mapping the latitude and longitude coordinates of the birth place or current address (1 km resolution) to Middle Layer Super Output Area (MSOA) regions (Figure 1). MSOA regions are defined as a set of adjacent output areas designed to have comparable population sizes and to be: “as socially homogeneous as possible based on tenure of household and dwelling type”.^20^ We ran separate analyses with dummy variables based on birthplace, and dummy variables based on current address. We only included regions with ≥ 100 UK Biobank participants, which resulted in 859 regions for the birth place analyses and 1,959 regions for the current address analyses, and a total of 271,635 participants with both birth place and current address available (Figure 1). As we address more fully in the methods, the current address measure is likely to be more precise than the birthplace measure. In order to quantify the impact of controlling for geographic confounding, we compared the GWAS results controlling for region with the results of conventional GWASs not corrected for region, obtained from the same selection of individuals. The impact of controlling for geographic confounding was investigated by computing the magnitude and significance of the change in SNP-based heritability and of the change in genetic correlation with two indicators of SES, namely educational attainment and household income.

**Figure 1:**
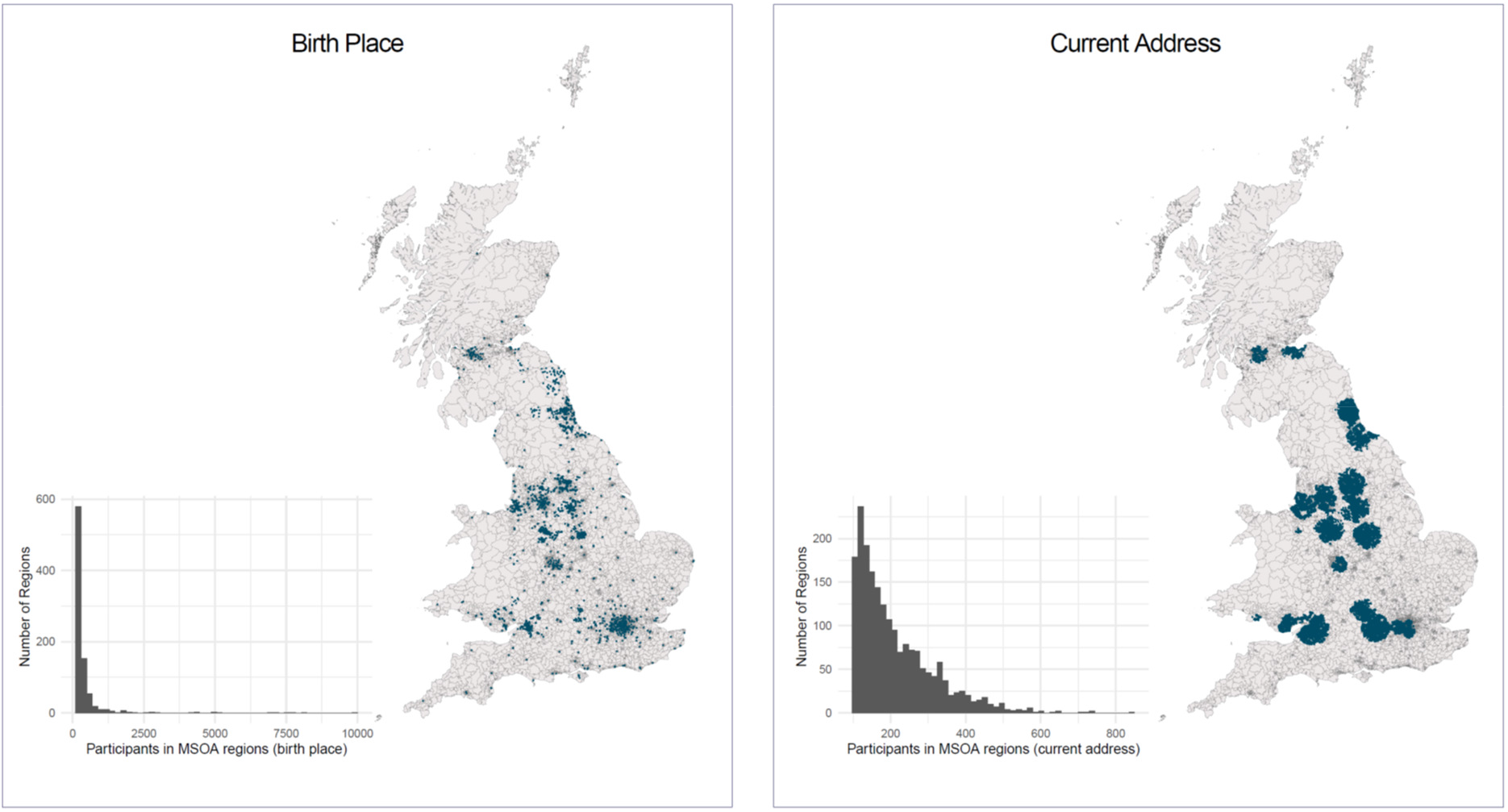
MSOA regions and the number of included UK Biobank participants for birth place and current address. The histograms show the distributions of the 271,635 UK Biobank participants the MSOA regions for birth place (859 regions, left) and for current address (1,959 regions, right).

### SNP-based heritability

Before controlling for geographic confounding, the SNP-based heritabilities of the 56 traits ranged from .02 to .41 (as estimated using LD score regression^10^, based on ~1.2 million SNPs), with physical traits showing higher heritability estimates than behavioral outcomes (Figure 2). The average SNP-based heritability per category was: anthropomorphic = .22, cardiovascular = .17, cognition & SES = .10, depression = .07, physical health = .07, reproduction = .11, sleep = .07, social = .04, substance use = .05, and other behavioral traits = .05. Figure 2 shows the change in individual SNP-heritabilities before and after controlling for geographic fixed effects. All corrected and uncorrected heritability estimates and the significance of their change can be found in *Supplementary_File_h2.xlsx*.

**Figure 2:**
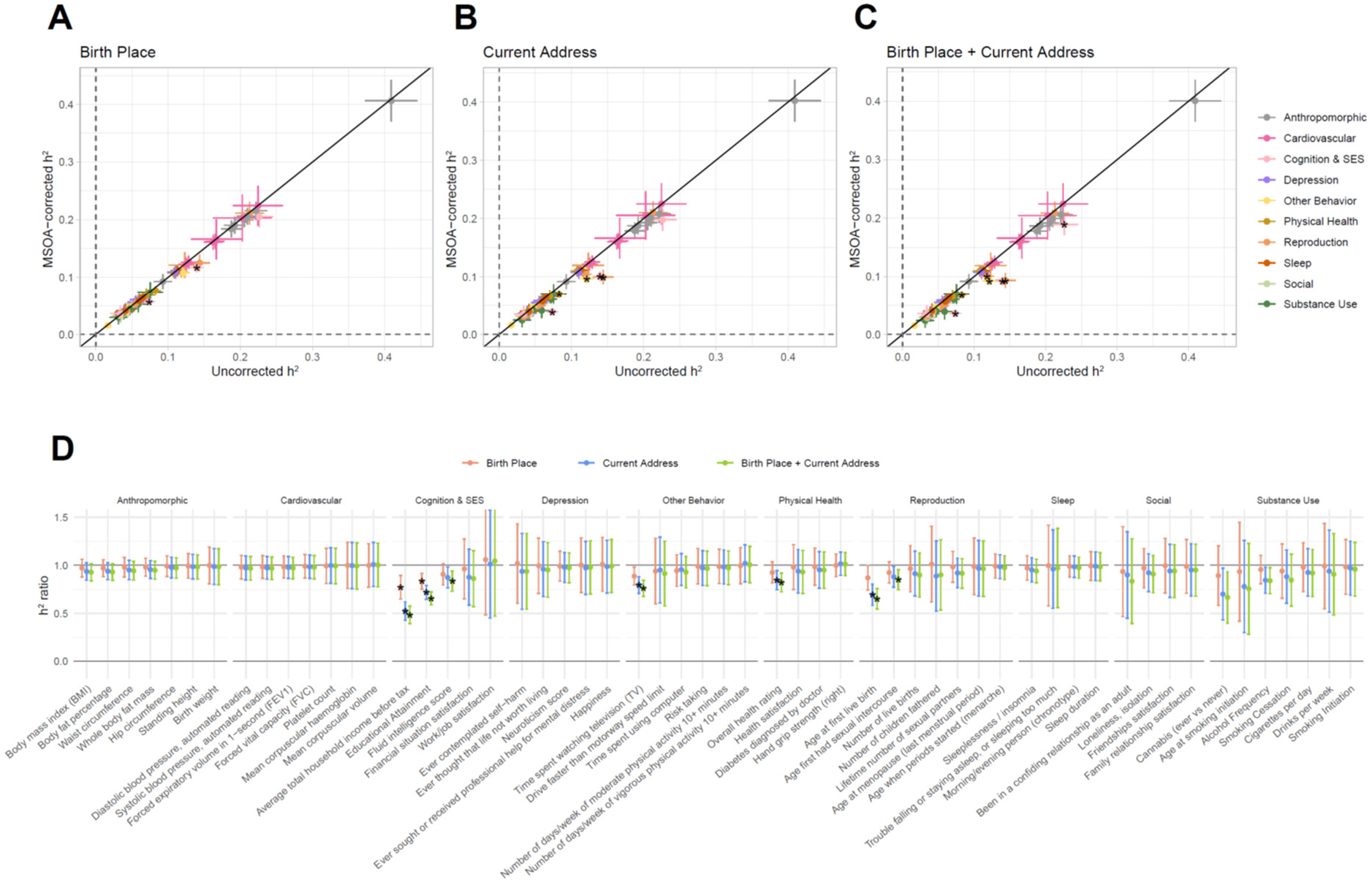
SNP-based heritabilities of 56 complex traits, corrected and uncorrected for MSOA region. Panels A-C show the estimated SNP-based heritabilities after controlling for MSOA regions based on birth place and/or current address. Panel D shows the ratio of the decrease, computed by dividing the corrected by the uncorrected SNP-based heritability estimate. The black stars indicate FDR-corrected p-values < 0.05, indicating significant changes in SNP-based heritability.

In GWASs that controlled for geographic fixed effects based on birth place (i.e., passive gene-environment correlations), the SNP-based heritability showed a significant decrease for two traits, namely *educational attainment* (from 14% to 12%) and *household income* (from 7% to 6%). When correcting for current address (i.e., a mix of passive and active gene-environment correlations) the SNP-based heritability significantly decreased for five traits, namely *household income* (from 7% to 4%), *educational attainment* (from 14% to 10%), *age at first birth* (from 14% to 10%), *time spent watching television* (from 12% to 10%), and *overall health* (from 8% to 7%). When correcting for both birth place and current address simultaneously, the SNP-based heritability significantly decreased for seven traits, namely *household income* (from 7% to 4%), *educational attainment* (from 14% to 9%), *age at first birth* (from 14% to 9%), *time spent watching television* (from 12% to 9%), *overall health* (from 8% to 7%), *age at first sexual intercourse* (from 12% to 10%), and *fluid intelligence* (from 23% to 19%).

### Genetic Correlations

Genes associated with socio-economic success can have an influence on the neighborhood that people can afford to live in, and thus on the quality of people’s living environment. The environmental exposures that differ between neighborhoods and regions can have effects on a wide range of physical and mental health outcomes, which can cause genes that are associated with socio-economic success to also become associated with these physical and mental health outcomes. We investigated whether controlling for regional differences decreases genetic correlations with SES by comparing genetic correlations before and after correcting for geographic region. The genetic correlations were computed between the complex traits and educational attainment and household income using LD score regression^21^ on 1.2 million SNPs, and the significance of the change in genetic correlations was tested in Genomic SEM^22^ accounting for the dependence between the various GWASs, which all rely on the same sample. Figure 3 shows the changes in genetic correlations and their significance. All uncorrected and corrected genetic correlations with educational attainment and household income can be found in Supplementary Figures 1 & 2 and in *Supplementary_File.xlsx*.

**Figure 3:**
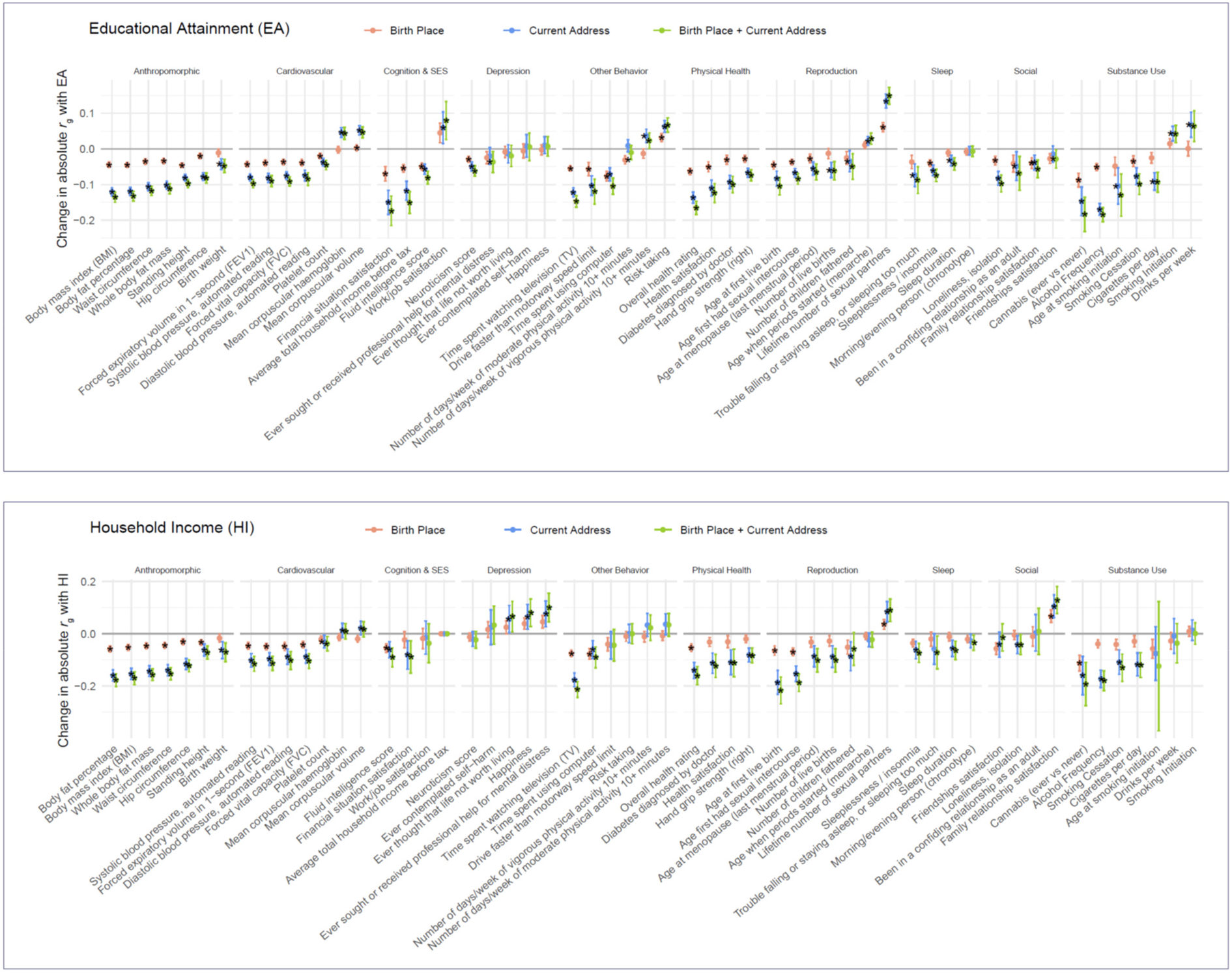
The change in absolute genetic correlations with educational attainment (EA, top) and household income (HI, bottom). The genetic correlations were computed with LDSC regression. We display the change in absolute genetic correlation in order to visualize the change in the strength of the genetic relationships with EA/HI (the directions of the genetic correlations vary between traits and are displayed in Supplementary Figures 1 & 2). The black stars indicate FDR-corrected *p*-values < 0.05, indicating significant changes in genetic correlation.

#### Genetic correlations with educational attainment

For 35 out of 55 complex traits we estimated a modest but significant change (FDR corrected) in genetic correlation with educational attainment when including fixed effects for MSOA region of birth place, 48 out of 55 when including fixed effects current address, and 50 out of 55 when including fixed effects for both birth place and current address. Most of the traits tested showed a significantly *weaker* genetic correlation with educational attainment (32 for birth place, 39 for current address, and 41 for both birth place and current address). When controlling for birth place and current address simultaneously, the five most significant decreases were observed for *body fat percentage* (from −.33 to −.20, *p*_change_=1×10^−82^), *BMI* (from −.30 to −.17, *p*_change_=1×10^−80^), *time spent watching TV* (from =−.69 to =−.55, *p*_change_=5×10^−76^), *alcohol frequency* (from −.42 to −.24, *p*_change_=9×10^−76^), and *overall health* (from −.49 to −.33, *p*_change_=2×10^−68^).

There was a relatively small portion of traits that showed significantly *stronger* genetic correlations with educational attainment after controlling for geographic region. When controlling for birth place, the genetic correlation with education increased for *lifetime number of sexual partners*, *risk taking*, and, *mean corpuscular volume*. When controlling for either current address or for both current address and birth place, the same three traits showed significantly stronger genetic correlations with educational attainment, as well as an additional six traits, namely *drinks per week*, *smoking initiation*, *age at menarche*, *mean corpuscular haemoglobin*, *work/job satisfaction*, and *days per week of vigorous physical activity 10*+ *minutes*. This could mean that regional differences in SES masked the genetic correlations between these traits and educational attainment, but could potentially also be a result of collider bias (see Discussion & Methods).

#### Genetic correlations with household income

For 19 out of 55 traits, we observed a significant change in their genetic correlation with household income when including fixed effects for MSOA region of birth place, 40 out of 55 when including fixed effects for current address, and 42 out of 55 when including fixed effects for both birth place and current address. Most of the traits tested showed a significantly *weaker* genetic correlation with household income (17 for birth place, 33 for current address, and 35 for both birth place and current address). For birth place + current address, the five most significant decreases were observed for *body fat percentage percentage* (from −.32 to −.14, *p*_change_=2×10^− 46^), *time spent watching TV percentage* (from −.62 to −.41, *p*_change_=2×10^−43^), *whole-body fat mass percentage* (from −.25 to −.09, *p*_change_=2×10^−42^), *BMI* (from −.33 to −.16, *p*_change_=4×10^−41^), and *waist circumference percentage* (from −.29 to −.14, *p*_change_=6×10^−38^).

There was a relatively small portion of traits that showed significantly *stronger* genetic correlations with income after controlling for geographic region. When controlling for birth place, the genetic correlation with income increased for *lifetime number of sexual partners*, and, *family relationship satisfaction*. When controlling for either current address or for both current address and birth place, the same two traits showed significantly stronger genetic correlations with educational attainment, as well as an additional five traits, namely *ever thought that life is not worth living*, *ever sought or received professional help for mental distress*, *mean corpuscular volume*, *mean corpuscular haemoglobin*, and *happiness*. This could mean that regional differences in SES masked the genetic correlations between these traits and income, but could potentially also be a result of collider bias (see Discussion & Methods).

## Discussion

We found traces of geographic confounding in the GWAS signals of a wide range of physical and mental health, body composition, and emotional, cognitive, behavioral, and socio-economic outcomes. The environmental effects that differ between rich versus economically deprived regions become entangled in the GWAS signals of most complex traits, as reflected their heritability estimates and their genetic correlations with SES-related traits. One efficient way to control for SES-related gene-environment correlations is by conducting within family GWAS analyses.^23–25^ Within family analyses, however, do not allow us to identify and study the specific source of gene-environment correlations, where our analyses do. In addition, while family-datasets are growing, large sample sizes of genotyped families are harder to attain, which results in less powerful within-family GWASs with noisier estimates of genetic effects (Figure 4). We show here that confounding due to gene-environment correlations can extend beyond the family environment and that this confounding can be reduced in existing large-scale datasets of both related and unrelated individuals by controlling for the region that participants were born in or moved to.

**Figure 4:**
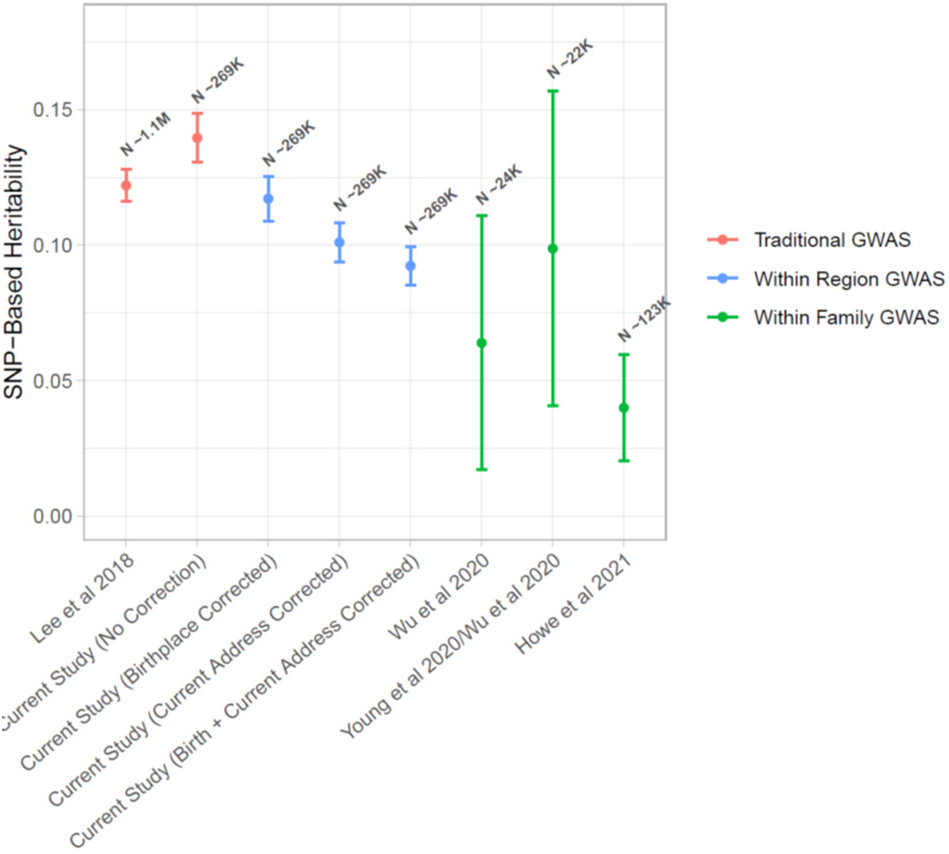
SNP-based heritability estimates of educational attainment (EA) under different GWAS designs.

After controlling for place of birth, the SNP-based heritability significantly decreased only for traits that directly reflect SES, namely educational attainment and income. Controlling for current address significantly decreased the heritability estimates for education and income more strongly, and significantly decreased the heritability for four additional traits, including overall health. Controlling for both birth place and current address simultaneously, resulted in two more traits with significantly reduced heritability estimates, including fluid intelligence. In contrast to the SNP-based heritability, the genetic correlation with SES-related outcomes (education and income) decreased significantly for many more traits investigated. The reduction in SES-related polygenic signal is more widespread and more substantial and significant when controlling for current address than for birth place, which suggests stronger confounding due to active gene-environment correlations (e.g., migration) than due to passive gene-environment correlations (e.g., rearing environment). An additional mechanism that could contribute to differences in heritability and genetic correlations when controlling for current address rather than birthplace is the passive wealth transfer from parent to child throughout life (e.g., financial support or inheritance).

The most significant reductions in genetic correlations with education and income were observed for traits related to BMI and body fat, suggesting that gene-environment correlations may especially come from exposures that are related to obesogenic environments and/or the ability or means to stay healthy and exercise. This may help partly explain why the polygenic score for BMI shows the strongest geographic clustering in Great Britain, after educational attainment and cognition.^15^ While the relationship between BMI and SES is positive in poorer countries, where food insecurity relates to a shortage of food in general, it is reversed in more developed countries, where food insecurity relates to a lack of access to healthy foods.^26^ Both the amount of fast-food restaurants and the diabetes rates in British neighborhoods are significantly correlated with regional differences in educational attainment, with more fast-food restaurants and higher diabetes rates in regions with lower average educational levels, but also with lower educational attainment polygenic scores.^15^ Behavioral traits that are related to body weight, namely time spent watching television, time spent using the computer, and alcohol intake, are also among the top five traits that showed the most significant decrease in their genetic correlation with educational attainment and income after controlling for geographic region. This suggests that, besides dietary options, there are behaviors that are correlated with regional socio-economic factors that affect regional differences in body weight and their associated genome-wide polygenic signals.

There are several limitations that have to be kept in mind when interpreting our results. Firstly, the UK Biobank is not a random sample of the general population, both in terms of participants’ phenotypes and with respect to their geographic locations. UK Biobank participants are, on average, higher educated and more healthy than the general population, and are more often from regions that are less economically deprived.^27,28^ They have been sampled to live within a radius of 35 km of one of the 22 UK Biobank assessment centers, so while for birth place there is wider geographic coverage, there is narrower geographic coverage for current address. It is unclear to what extent this ascertainment bias contributes to the geographic confounding in the GWAS and to our ability to control for it. Secondly, the geographic locations we use are not precise: for the purposes of anonymity of the participants, their birth and current home location have been rounded to 1 km, which we then mapped to nearest MSOA region. While both birth place and current address are rounded, the locations for current address are likely to be more precise than those of the birth place (see Methods). More accurate locations may improve our ability to control for geographic confounding. Thirdly, while boundaries of MSOA regions have been chosen to delineate socially homogeneous regions,^20^ the geographic location of the participants is only a relatively crude and temporally variable proxy for the social environment that underlies the gene-environment correlations we try to control for. Environmental circumstances start with the family environment, and expand to close and extended social circles, which include schools, peers, work environment, and thus may extend across multiple communities throughout life. It will be challenging to fully understand and account for the correlation between genes and environments, but there is room for improvement by genotyping family members, social circles, and collecting longitudinal information on the participants’ living environments. Finally, when correcting for geographic location, there is a risk of collider bias: if a genetic variant affects current address (for example through its effect on cognitive ability and therefore educational attainment), and the outcome of interest (for example substance use) also affects current address, then controlling for current address in the GWAS on substance use may induce collider bias, which biases the association between genetic variant and substance use, and biases the downstream estimates of heritability and genetic correlations.

To summarize, we detected and controlled for part of the confounding due to gene-environment correlations on a regional level. Our findings suggest that effects estimated in GWASs of many phenotypes are confounded, and that this confounding is not entirely attributable to processes that take place within a family as is implied by terms like “genetic nurture”, but also more broadly attributable to social and political processes that correlate to individuals’ genotypes. If GWASs are to remain central to the study of (non-communicable) disease epidemiology and biology, their designs, and conclusions, need to carefully reflect the reality of the social and geographic structure of society.

## Methods

### Participants

The participants of this study come from UK Biobank.^29,30^ UK Biobank has received ethical approval from the National Health Service North West Centre for Research Ethics Committee (reference: 11/NW/0382). A total 273,402 females and 229,134 males (N_total_ = 502,536) aged between 37 and 73 years old were recruited between 2006 and 2010 across 22 assessment centers throughout Great Britain. The sampling strategy was aimed to cover a variety of different settings providing socioeconomic and ethnic heterogeneity and urban–rural mix. The participants underwent a cognitive, health, and lifestyle assessments, provided blood, urine, and saliva samples, and have their health followed longitudinally.

### Genotypes and Quality Control (QC)

A total of 488,377 participants had their genome-wide SNPs genotyped on the UK BiLEVE array (N = 49,950) or the UK Biobank Axiom Array (N = 438,423). Genotypes were imputed using the Haplotype Reference Consortium (HRC) panel as a reference set (pre-imputation QC and imputation are described in more detail in Bycroft et al, 2018).^30^ We extracted SNPs from HapMap3 (1,345,801 SNPs) from the imputed dataset. In the pre-PCA QC on unrelated individuals, we filtered out SNPs with MAF < .01 and missingness > .05, leaving 1,252,123 SNPs. After filtering out individuals with non-European ancestry (see paragraph on *Ancestry & Principal Component Analysis* below), we repeated the SNP QC on unrelated Europeans (N = 312,927), filtering out SNPs with MAF < .01, missingness >.05 and HWE p < 10^−10^, leaving 1,246,531 SNPs. We then created a dataset of 1,246,531 QC-ed SNPs for 456,064 UK Biobank subjects of European ancestry.

### Ancestry & Principal Component Analysis

Ancestry was estimated using Principal Component Analysis (PCA). We first determined which participants had non-European Ancestry by projecting the UK Biobank participants onto the first two principal components (PCs) from the 2,504 participants of the 1000 Genomes Project, using HM3 SNPs with minor allele frequency (MAF) > 0.01 in both datasets. Participants from UKB were assigned to one of five super-populations from the 1000 Genomes project: European, African, East-Asian, South-Asian, or Admixed. Assignments for European, African, East-Asian, and South-Asian ancestries were based on > 0.9 posterior-probability of belonging to the 1000 Genomes reference cluster, with the remaining participants classified as Admixed. Posterior-probabilities were calculated under a bivariate Gaussian distribution, where this approach generalizes the k-means method to take account of the shape of the reference cluster. We used a uniform prior and calculated the vectors of means and 2×2 variance-covariance matrices for each super-population. A total of 456,064 subjects were identified to have a European ancestry. In order to capture ancestry differences within the British population, a PCA was then conducted on these 456,064 individuals of European ancestry. When trying to capture ancestry differences in homogenous populations, genotypes should be pruned for LD and long-range LD regions removed.^31^ The LD pruned (r^2^ < .2) UKB dataset without long-range LD regions consisted of 131,426 genotyped SNPs. The PCA to construct British ancestry-informative PCs was conducted on this SNP set for unrelated individuals using flashPCA v2.^32^ PC SNP loadings were used to project the complete set of European individuals onto the PCs. PCs that reflect ancestry differences are expected to cluster geographically, which we investigated by computing their Moran’s *I*: out of the top 100 PCs, 78 PCs showed significant geographic clustering after Bonferroni correction, and 94 PCs showed a *p*-value < .05 (see Abdellaoui et al, 2019, on details on evaluating the geographic clustering of the PCs using Moran’s *I*)^15^.

### Genetic Relatedness Matrix

We created genetic relatedness matrices (GRMs) to include in our LMM GWAS in order to better control for population stratification and to control for cryptic relatedness. The GRMs contain genetic relationships between all individuals based on a slightly LD pruned HapMap 3 SNP set (LD-pruning parameters used in PLINK: window size = 1000 variant count, step size = 100, r^2^-cutoff = 0.9 and MAF > 0.01, resulting in 575,293 SNPs). The GRMs were computed using GCTA^33^ on individuals of European descent. We created a sparse GRM, containing only the relationships of related individuals (cut-off = .05, resulting in 179,609 relationships), as one of the major goals of including the GRM in our LMM GWASs was to control for the presence of closely related subjects (cousins, siblings, parent-offspring).

### Phenotypic & Geographic Measures

#### Phenotypes

The selection of phenotypic outcomes was based on the relevance of the metric to broad mental and physical health outcomes, resulting in the selection of 56 complex traits encompassing domains of anthropomorphic traits, cardiovascular outcomes, cognition, SES, depressive symptoms, sedentary behavior, reproductive behavior, risk taking behavior, physical activity, self-reported overall health, sleep, social connection, and substance use (see Supplementary Table 1). All traits were analyzed as provided by UK Biobank, except for the substance use phenotypes, which were defined according to the GSCAN GWASs in Liu et al (2019)^34^, and educational attainment, which was transformed to years of education as defined according to the ISCED coding as analyzed in Lee et al (2018)^35^.

#### Geographic measures

Birth place location was based on the coordinates in the UK Biobank fields 129 (latitude) and 130 (longitude). The UK Biobank verbal interview includes a procedure to ascertain participant birthplace described as follows: “*The interviewer is provided with a tree structure that lists place names and counties in England, Wales and Scotland. They were instructed to enter at least the first 3 letters of the town/village/place that the participant provides. If there are too many matches to the first 3 letters, no place names appear, and more letters need to be entered. If there is more than one listing of the relevant place name, then they were asked to choose the one with correct district or county. In order to narrow down the search, the interviewer may also need to type in the district or county to find a match. If they cannot find the place name, they were instructed to use the Enter Other facility to enter free text to state the town, county (or district) and country. If the participant does not know, there is the option to enter Unknown. Place of birth in the UK is then converted into North and East co-ordinates.*” It is not clear whether birth place refers to the residence at birth or the place of the hospital of birth. Current address location was based on the coordinates in the UK Biobank fields 22702 (longitude) and 22703 (latitude). Deriving these coordinates is described by UK Biobank as follows: “*Where the full address is present and is verified to be valid, software package DataPlus (provided in the QuickAddress Batch packet), is used to transform the address data into the grid coordinates. Where only the postcode is present, the grid coordinates were generated with the aid of on-line mapping tools: doogal [https://www.doogal.co.uk/] and uk-postcodes [https://www.uk-postcodes.com]. Further details of this process are available from relevant websites.*“ The participants’ coordinates, which were rounded to 1km, were mapped to the nearest MSOA region using a shape file obtained from the InFuse website, which is part of the UK Data Service Census Support.^36^ The R-packages sp (v1.4-4) and rgdal (v1.5-18) were used to merge the spatial data from the MSOA shapefile.^37–39^

#### Genetic Association Analyses

We performed four GWASs per trait. First, we performed GWASs using two regression models. Model 1 is as follows:

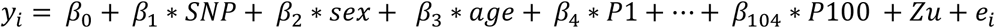

Where *P1* to *P100* represent ancestry-informative PCs derived as described above, *u* is a vector of random effects, and *Z* is a random effects design matrix and a random effects model as described in Jiang et al (2019)^40^, which is used to account for the presence of closely related subjects and an extra guard against population stratification. Model 2 is as follows:

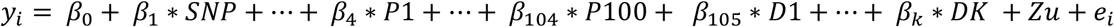

The second equation omits *sex* and *age* for brevity and includes dummy variables *D1* thru *DK* for all but 1 MSOA region of birth. The following DAG describes the suspected causal structure of the data we model:

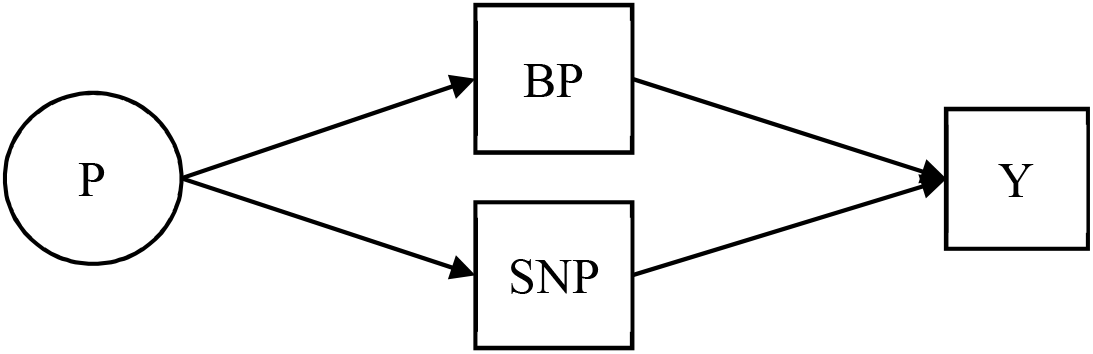

Where *P* represents the previous generation (parents), *BP* is birth place, *SNP* represents the genes carried by the individual, and *Y* the phenotypic outcome. Along similar lines, a GWAS was performed using dummy variables based on current address instead of birth place, and an analysis where dummy variables for both birthplace and current address were included. The plausible causal models in the GWASs that correct for current address are more complicated. We offer 3 DAGs which we feel are abstractions of potential causal processes that relate confounding parental influences (P), genotype (SNP), current address (CA), and outcome (Y). We include these DAGs and their descriptions, because they are useful to have in mind when evaluating our results. In practice we expect that a mix of these processes, or more complex processes altogether, underpin the relation between genotype and outcome.

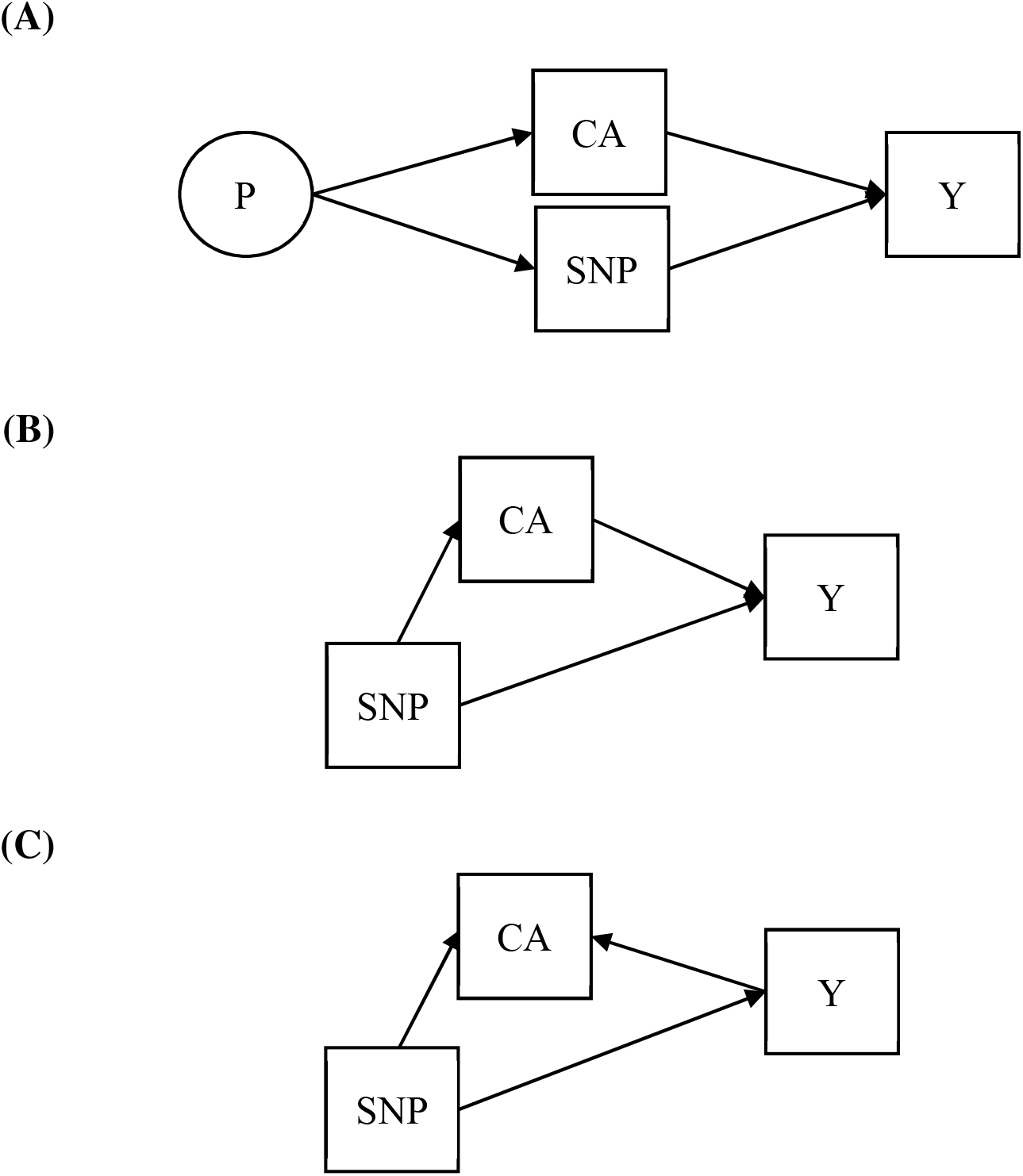

The first causal process (A) we suspect plays out is a process where a confounder, like intergenerational transfer of wealth through inheritance or financial support during college (or lack thereof), is correlated to the parental genome and therefore the offspring’s genome (*SNP*) as well as the address of the adult participant (*CA*), and controlling for address ensures the regression of *Y* on *SNP* is no longer confounded by *P* (similar to the process that underlies controlling for birth place).

The second causal process (B) is one where the genotype (*SNP*) mediated by traits like cognitive ability and mental health affects people’s ability to attain a higher education and/or be upwardly socially mobile, and so influences which environment people can afford to live in, which in turn influences the outcome (*Y*) of interest. Here, if we control for current address, we test for a more “direct” effect, not mediated by environmental exposures, of genotype (*SNP*) on outcome (*Y*). The (probable) presence of mediation would mean that controlling for current address is likely to lead to qualitatively changes in the genetic effect estimates to a different extent than when controlling for birthplace. However, even in the absence of mediation, we could expect differences herein, as current address appears to be measured more precisely.

Finally, as depicted in causal model C, there is the risk that the genotype (*SNP*), through its effect on other traits (e.g., cognitive ability), influences current address (*CA*), while the outcome (*Y*) (e.g., substance use) also influences current address. In this case, conditioning on current address when doing a GWAS on, e.g., substance use would potentially induce collider bias, which is undesirable. In practice we do not know which of the causal processes, or which mix of causal processes, underlies the data. The effects of correcting for current address are therefore more complex to interpret.

### SNP-Based Heritability and Genetic Correlation

Heritability and genetic correlation were estimated using LD Score regression^10,41^ and Genomic SEM^22^ version 0.0.3. The genetic correlations are based on the estimated slope from the regression of the product of z-scores from two GWASs on the LD score and represents the genetic covariation between two traits based on all polygenic effects captured by the included SNPs. The genome-wide LD information used were based on European populations from the HapMap 3 reference panel.^42,43^ All LD score regression analyses included the ~1,3 million genome-wide HapMap SNPs used in the original LD score regression studies.^42,43^ The standard error of the difference between the heritability and genetic correlations based on the different specifications cannot easily be estimated directly, because the GWASs for which we want to obtain the differences are based on the exact same sample, and their standard errors are therefore highly dependent. Therefore, we estimated the standard error of the differences in heritability and genetic correlations with Genomic SEM, which allows us to account for the dependence between the estimates of the SNP-based heritability and genetic correlations.

## Supporting information

Supplementary_File_rgs.xlsx

## Acknowledgements

This study was conducted using UK Biobank resources under application number 40310. A.A. & K.J.H.V. are supported by the Foundation Volksbond Rotterdam. A.A. is also supported by ZonMw grant 849200011 from The Netherlands Organisation for Health Research and Development. M.G.N is supported by the National Institute Of Mental Health of the National Institutes of Health under Award Number R01MH120219, ZonMW grants 849200011 and 531003014 from The Netherlands Organisation for Health Research and Development, a VENI grant awarded by NWO (VI.Veni.191G.030), and is a Jacobs Foundation Fellow.

## Code availability

Code will be made available on github: https://github.com/MichelNivard/region_FE_GWAS

## Supplementary Figures & Tables

**Supplementary Figure 1:**
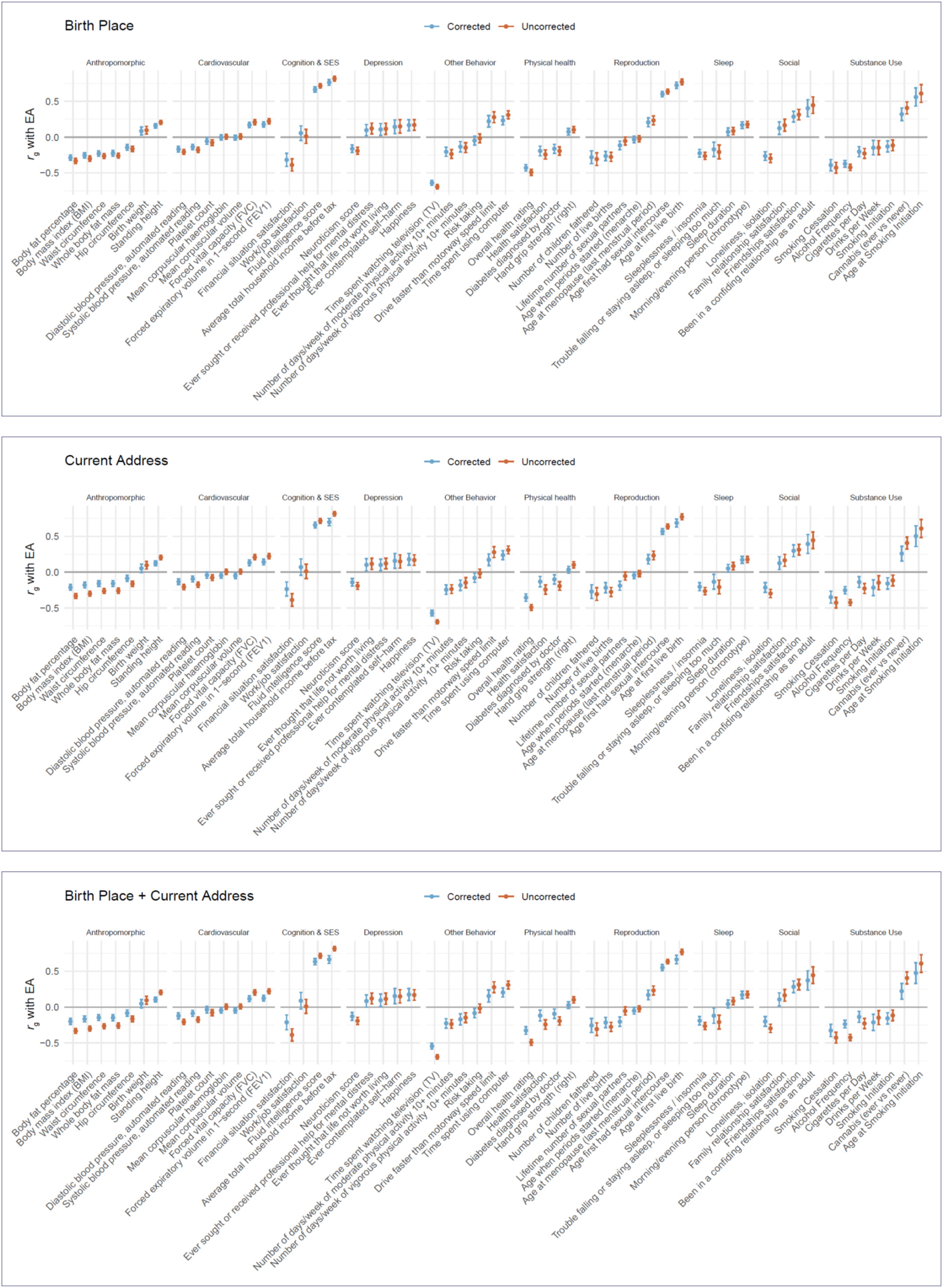
Genetic correlations (*r*_g_) with educational attainment (EA) as computed with LDSC regression, before and after controlling for MSOA region of birth place (top), current address (middle), or birth place + current address (bottom).

**Supplementary Figure 2:**
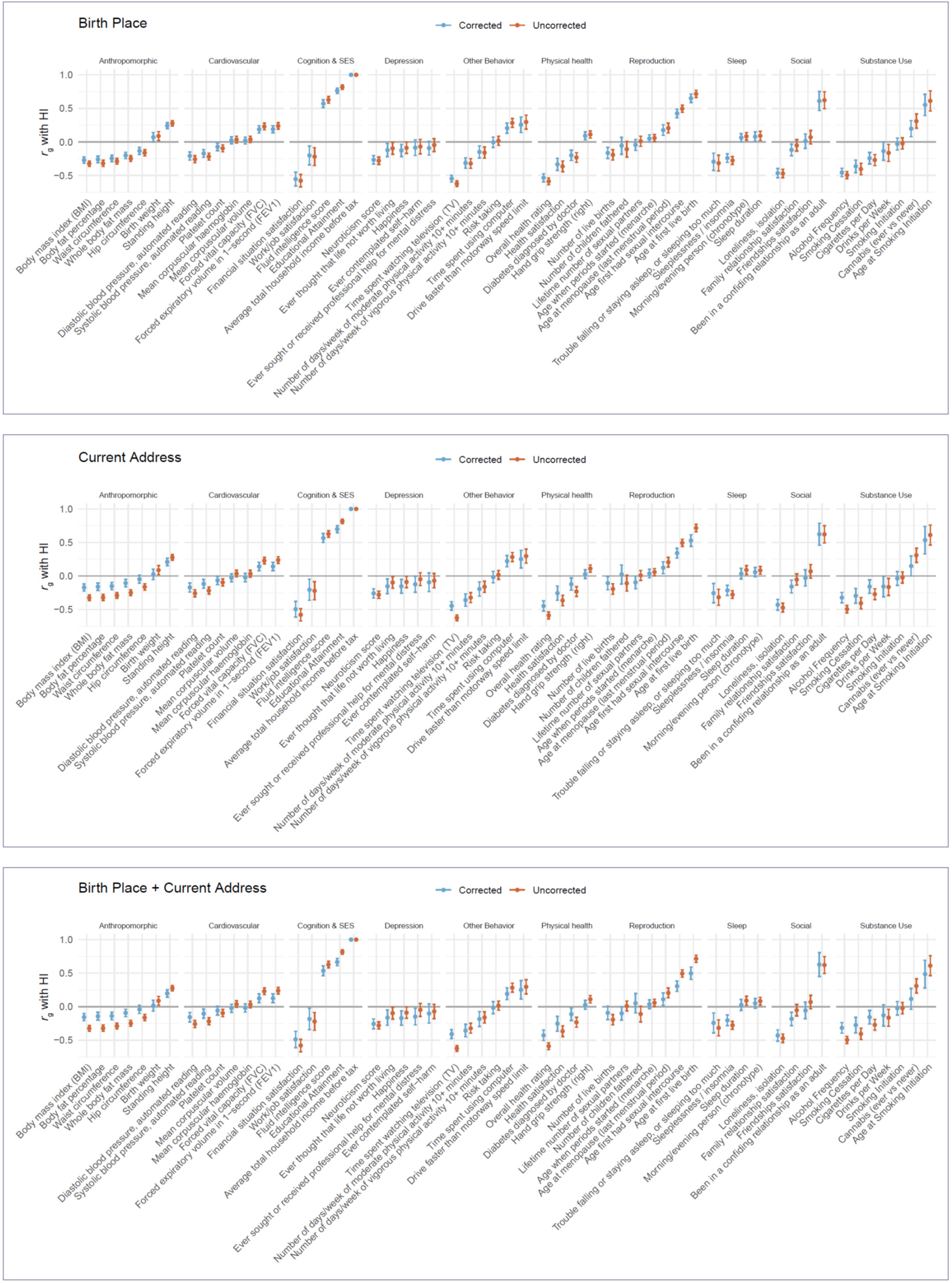
Genetic correlations (*r*_g_) with household income (HI) as computed with LDSC regression, before and after controlling for MSOA region of birth place (top), current address (middle), or birth place + current address (bottom)..

**Supplementary Table 1:**
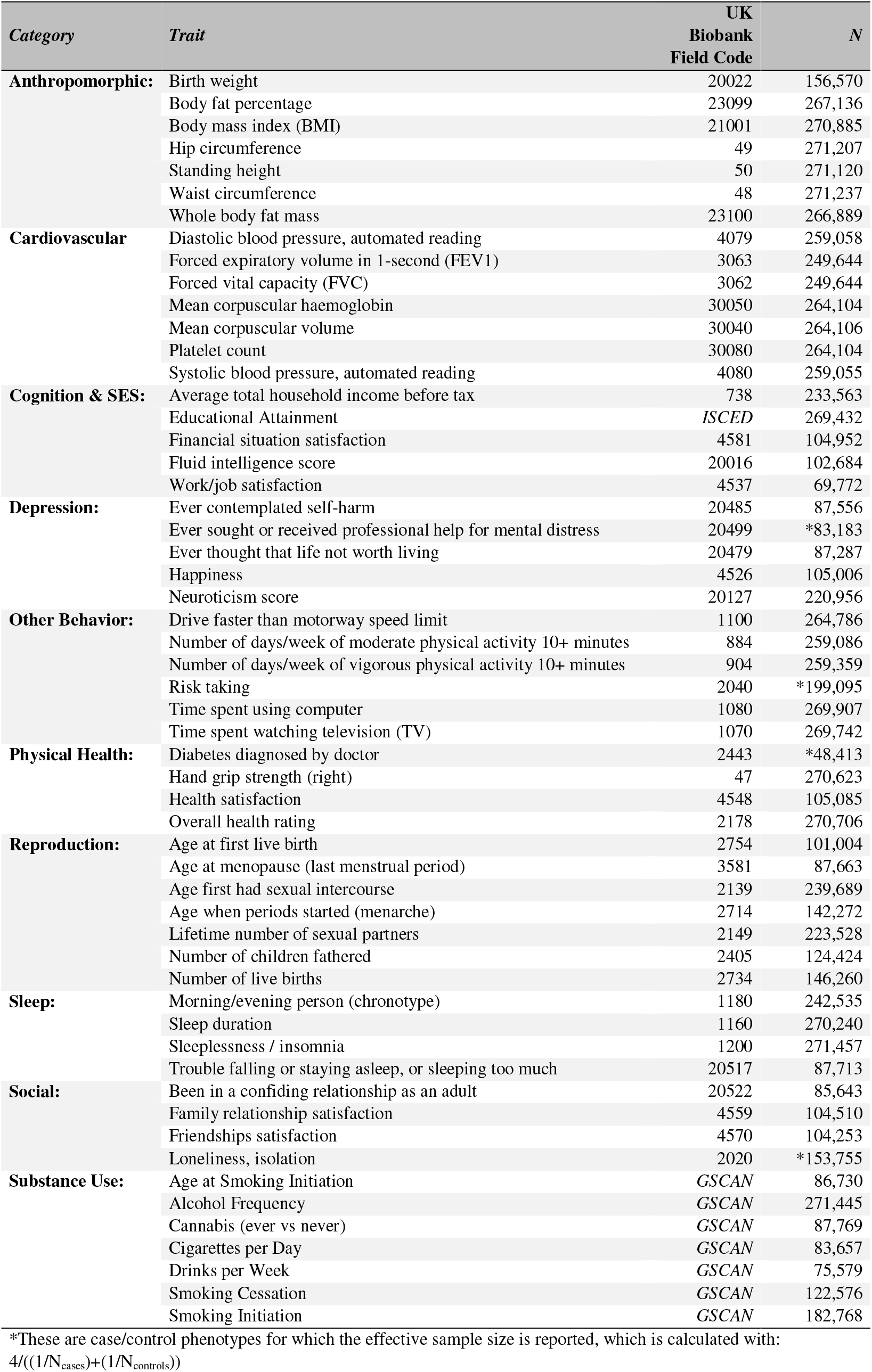
The 56 complex traits and the (effective) sample sizes of their GWAS analyses.

